## Supplemental Table 2 for "Proteome-wide antigenic profiling in Ugandan cohorts identifies associations between age, exposure intensity, and responses to repeat-containing antigens in *Plasmodium falciparum*"

**Supplementary table 2 – Top Proteins with highest seropositivity (>38%) in the cohort**

| **Known targets of protective antibodies in humans or animal models** | | **No evidence as targets of protective antibodies** | |
| --- | --- | --- | --- |
| *Characterized in high-resolution in this study* | *Previously*  *well-characterized* | *Characterized in high-resolution in this study* | *Previously*  *well-characterized* |
| PHISTc(PF3D7_0801000) ^1^  MSP10^2^  Rh2a ^3,4^  Rh2b ^3,4^  EMP3 ^5^  RON2^6^  DBLMSP^7^ | Ag332 ^8–10^  LSA3^11^  STARP^12^  GLURP^13–15^  MSP4^16,17^  MSP2^18–20^  EBA175 ^21,22^  KAHRP^23^  MSP1^24^  PF11-1^25^  RIFIN ^26,27^  STEVOR ^28^  PfEMP1^29,30^ | SURFINs  REX1  SBP1  TREP  PF3D7_0424700(FIKK Ser/Thr Kinase)  PF3D7_0511500 (RNA pseudouridylate synthase)  PTP3  RON4  LISP2  PTP4  CRMP4  PF3D7_0628100  PF3D7_1023100(putative dyenin heavy chain)  PF3D7_0907200(putative HECT domain-containing protein 1)  PFIT_1002300  PFIT_0522600  CRA(EXP1) | FIRA ^31^  MESA ^32,33^  LSA1 (Protective T cell responses - ^34^) |
